# Macrophage Cholesterol Homeostasis Underpins the Role of Sertraline as an Adjunctive Agent in Tuberculosis Therapy

**DOI:** 10.1101/2025.07.18.665474

**Authors:** Kanika Bisht, Riya Sahu, Sneha Vasudevan, Simran Jit, Nikita Bhor, Jitendra Kumar, Mahima Madan, Deepthi Shankaran, Sherry Bhalla, Nagasuma Chandra, Vivek Rao

## Abstract

Cellular metabolism plays a deterministic role in the macrophage responses to intracellular pathogens such as *Mycobacterium tuberculosis* (Mtb) and infection control. Here, we demonstrate that the significant rewiring of the macrophage cholesterol metabolism by sertraline (SRT), an FDA-approved antidepressant, effectively enhances bacterial control. SRT, by virtue of its potent cationic amphiphilicity leads to the accumulation of lysosomal cholesterol in macrophages and activating the transcription factor SREBP2 and enhancing cholesterol biosynthesis. By specific gene silencing and biochemical inhibition assays, we further demonstrate that this metabolic reprogramming promotes lysosomal membrane permeabilization, mitochondrial reactive oxygen species (ROS) generation, heightened IL-1β release, collectively enhancing macrophage bactericidal activity. Our results thus highlight a previously underappreciated link between cholesterol homeostasis and inflammasome activation, which contributes to improved Mtb clearance in the presence of adjunctive sertraline therapy.

**Statement of significance:** Tuberculosis remains a leading cause of global mortality, and treatment success is limited due to a long treatment regimen. This study identifies macrophage cholesterol homeostasis as a central pathway to enhance anti-tubercular therapy. We show that Sertraline, an FDA-approved antidepressant, a cationic amphiphilic drug (CAD), reprograms cholesterol trafficking through NPC1 inhibition. This drives the consequential processing of the master regulator SREBP2, accompanied by lysosomal membrane permeabilization. Collectively, these lead to mitochondrial ROS production and activation of NLRP3 signalling, boosting inflammasome-mediated bacterial clearance. By establishing cholesterol sequestration as the mechanistic basis of host-directed activity of sertraline and distinguishing it from ineffective CADs, our work provides a mechanistic framework for developing cholesterol-modulating host-directed adjuncts to improve TB treatment outcomes.

**Graphical abstract: SRT induced lysosomal cholesterol accumulation results in enhanced bacterial control via inflammasome activation:** 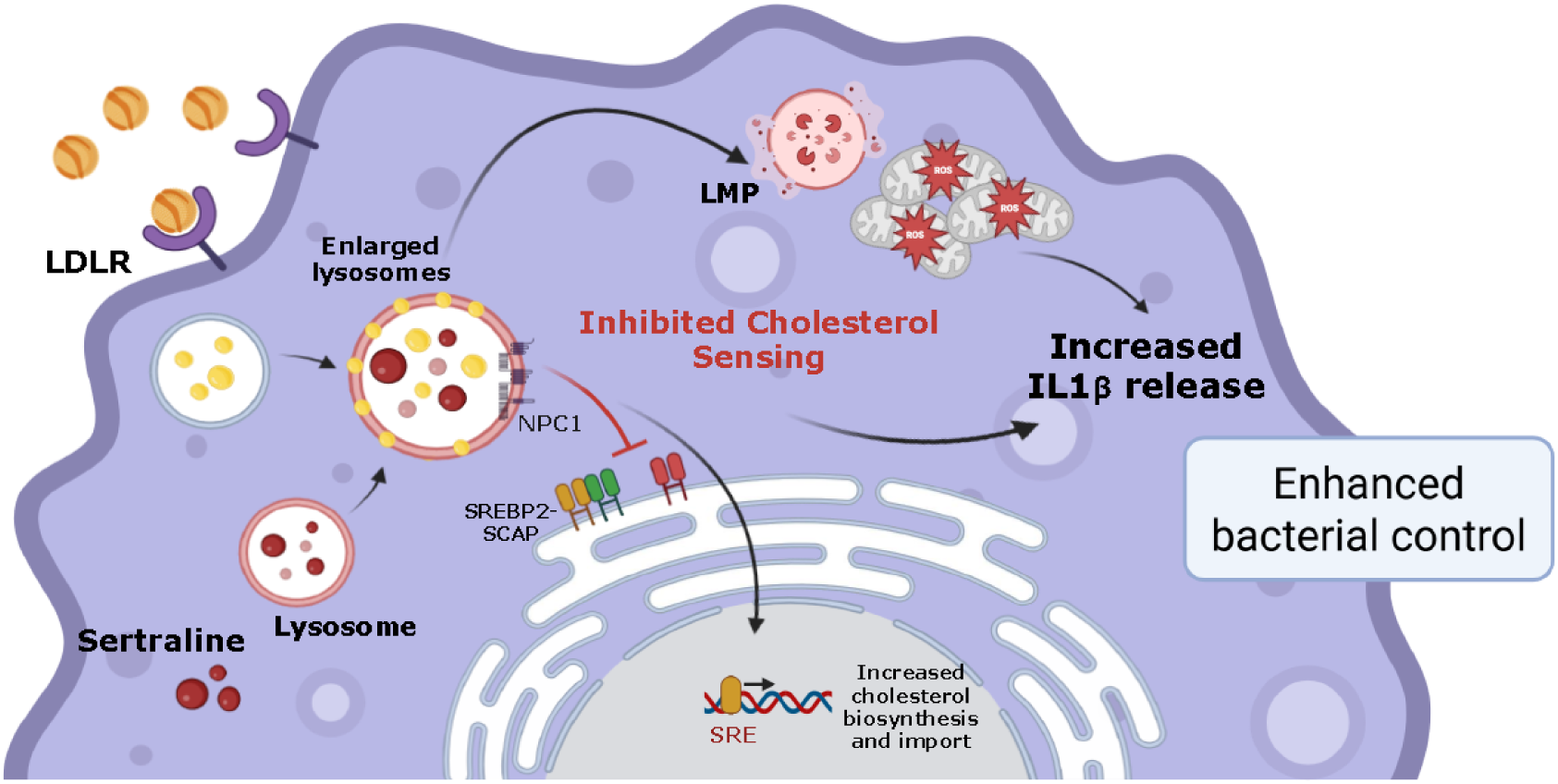

## Introduction

Tuberculosis (TB), caused by *Mycobacterium tuberculosis* (Mtb), remains the leading cause of death from a single infectious agent globally, outpacing all other bacterial and viral pathogens (1–3). The multiple initiatives to curb the infection have not been successful enough to eradicate the disease, instead have led to an increase in drug resistance in the population (4, 5). These limitations have spurred an interest in therapeutic strategies designed to aid the host in controlling infection. Such host-directed therapies (HDTs) have relied on harnessing the cellular physiological pathways to enhance antimicrobial properties (6–8). We have previously shown that sertraline (SRT), a selective serotonin reuptake inhibitor (SSRI), can act as a potent adjunct to standard anti-TB therapy (ATT), accelerating bacterial clearance, improving tissue resolution, and enhancing host survival in preclinical infection models (9). These properties are associated with the restriction of type I IFN responses via the activation of the host inflammasome pathway, thus defining the therapeutic potential of manipulating host immunity for infection control (10).

Recent findings strongly advocate for the influence of host sterol metabolism in regulating the inflammatory capacity of innate immune cells (11–13). More importantly, oxysterols have emerged as promoters of the pro-inflammatory capacity of macrophages (14, 15). Recently, SREBP2, the master regulator of cellular cholesterol metabolism has been associated with the direct activation of the host cell inflammasomes via the NLRP3 axis (16).

Here, we demonstrate that the ability of SRT to adjunctively augment intracellular bacterial control with frontline TB drugs is contingent on the significant alteration of macrophage cholesterol homeostasis. We demonstrate that treatment with SRT leads to a significant increase in intracellular cholesterol levels in Mtb-infected macrophages. We sought to investigate the mechanistic basis and consequence of this increase on bacterial growth control by employing chemical modulation and gene silencing, and demonstrate that the SRT induces significant cholesterol buildup in the macrophage lysosomes, resulting in distinct membrane permeabilization (LMP) and mitochondrial reactive oxygen species (ROS) production. We thus highlight a novel connection between cellular sterol homeostasis and the inflammasome mediated IL1β release in Mtb-infected macrophages. Taken together, our findings provide a strong mechanistic link between the cellular metabolic machinery (cholesterol metabolism) and antimicrobial efficacy. This work offers both conceptual and translational insights into leveraging host metabolism as a host-directed therapeutic strategy to improve TB treatment outcomes.

## Results

### Treatment with sertraline increases cellular cholesterol in macrophages

In an attempt to decipher the underlying mechanism for sertraline- mediated increase of antibiotic efficacy in Mtb infections, global expression profiles of Mtb-infected THP1 macrophages with a combination of isoniazid (H) and rifampicin (R) with (SRT) were compared with HR-treated cells. A total of 160 genes of cholesterol homeostasis were differentially upregulated on treatment with HRS (HR with SRT) when compared to HR treatment (Figure 1A). Interestingly, cholesterol homeostasis emerged as the most significantly enhanced metabolic pathway along with an increase in the mTORC1 signaling complex in macrophages treated with HRS (Figure 1B). In fact, HRS-treated macrophages displayed a significant increase in the expression of genes involved in cholesterol biosynthesis (Figure 1C) and import, with the markedly lower expression of sterol export genes (Figure 1D). A similar pattern was reflected in SRT or HRS-treated macrophages by qRT-PCR. While infection or treatment with HR failed to alter the expression of genes involved in cholesterol metabolism, genes involved in biosynthesis and import were 3-5-fold higher with treatment with HRS or SRT at 24 h of infection (Figure 1E). This increment in gene expression reflected as an SRT-dependent 1.2-1.5-fold increase in intracellular cholesterol (Figure 1F). To specifically inhibit cholesterol biosynthesis, we employed a pharmacological inhibitor of *HMGCR*, simvastatin. Contrary to our expectations of diminishing antibiotic efficacy, the addition of simvastatin significantly enhanced the control of Mtb by antibiotics by 3-5-fold in comparison to the cells treated with HR or HRS alone (Figure 1G). Previous reports have suggested that statins, in turn, can enhance expression of genes in cholesterol biosynthesis (17), we also tested for the gene expression following treatment with statins. Treatment with simvastatin induced the expression of the genes of the biosynthetic pathway as well as genes of cholesterol import. Also, a decrease in the cholesterol export genes has been observed with simvastatin treatment (Figure 1H). This suggested that the increased involvement of the pathway may be crucial in enhancing bacterial control by SRT.

**Figure 1:**
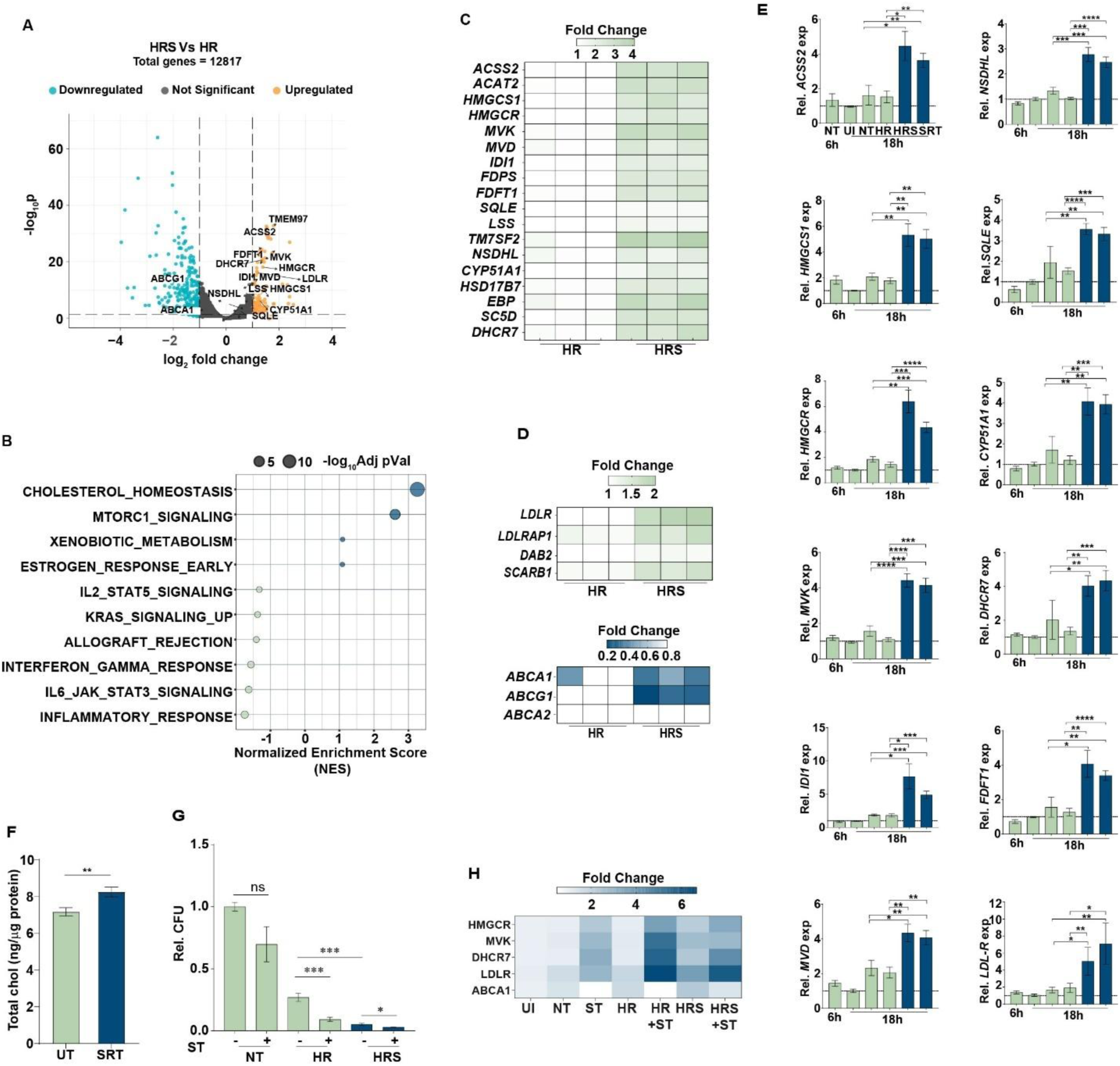
Global transcriptomics of Mtb-infected macrophages following treatment with sertraline. A) A comparison of the differentially expressed genes in Mtb-infected macrophages following an 18 h treatment with HR or HRS is shown as a volcano plot. Y- and X-axes denote the -log10 P values and log2 fold change, respectively. The upregulated genes with ≥ 2 fold change with p-value of ≤ 0.01 are highlighted in orange, while the downregulated genes with ≤ -2 fold change with p-value of ≤ 0.01 are in blue. B) GSEA analysis of the top significantly enriched gene sets in HRS vs HR is represented as a bubble plot. Gene sets that are significantly (P-value ≤ 0.05, 0.01) enriched (Blue) or downregulated (green) are represented, with the X-axis denoting the normalized enrichment score. C-E) Gene expression patterns in HR and HRS-treated macrophages: Expression of genes involved in cholesterol biosynthesis (C) and cholesterol trafficking (D), normalised to the untreated controls in the RNA-seq data was validated by gene-specific qRT-PCR (E). The relative expression of specific genes relative to *GAPDH* is represented as fold change w.r.t uninfected control (UI) at 6h + SEM of triplicate assays from 3 independent experiments (N=3). F) Cholesterol levels in THP1 macrophages after treatment with sertraline (SRT) for 24 hours. Total free cholesterol was normalized to the amount of protein and is represented as mean ± SEM for triplicate assays of three independent experiments (N=3). G) Growth of Mtb in HR or HRS treated macrophages that were treated with 10µM simvastatin (ST) for 3 days. Bacterial numbers are represented as mean log10 CFU ± SEM from triplicate assays of three or four independent experiments (N = 3 or 4). H) Effect of simvastatin on the expression of genes for biosynthesis (HMGCR, MVK, DHCR7), import (LDLR), and export (ABCA1) of cholesterol in Mtb-infected macrophages, after 18 h of treatment. The relative expression of specific genes, normalized to GAPDH, is represented as the fold change ± SEM of triplicate assays from three independent experiments (N = 3). P values less than 0.05, 0.01, 0.005 are represented as *, ** and *** respectively.

### Lysosomal cholesterol accumulation is crucial for Mtb control in macrophages

Cholesterol levels in different subcellular locations are tightly regulated to maintain a steady state in cells. To test the effect of increased cellular cholesterol on intracellular distribution, macrophages were stained with the cholesterol specific dye- filipin and visualized by fluorescence microscopy. While filipin staining was primarily concentrated on the membranes of untreated (UT) cells with minimal staining within, treatment with SRT severely impacted intracellular sterol levels, with cells displaying significantly higher intracellular staining in addition to the membrane (Figure 2A).

**Figure 2:**
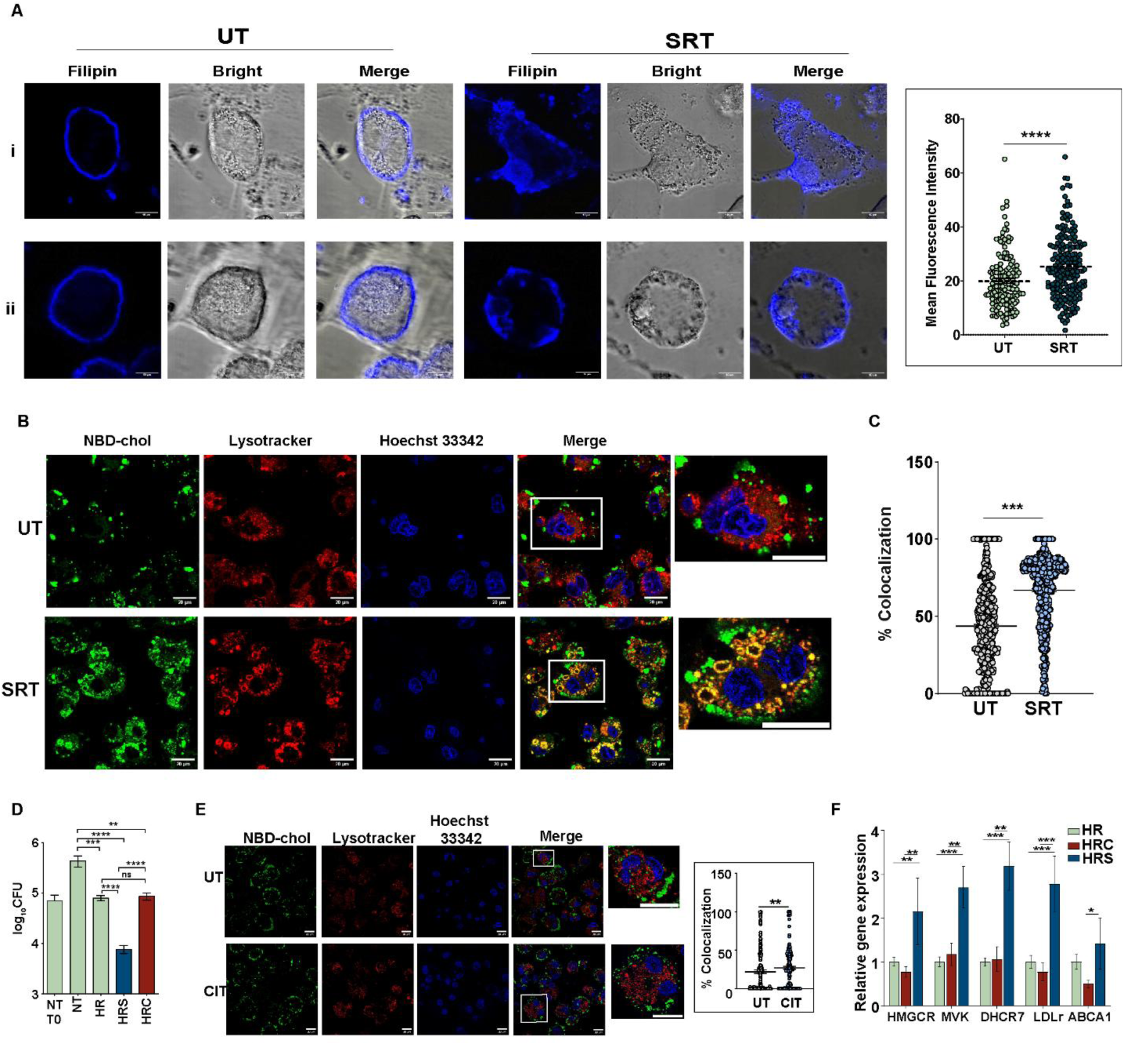
Sertraline treated macrophages accumulate lysosomal cholesterol. A-C) Cholesterol quantitation in THP1 macrophages after SRT treatment for 24 h. A) Representative confocal images of two independent cells (i and ii) stained for cholesterol with filipin and quantitated densitometrical (box). B) Representative confocal images of cells fed with NBD-cholesterol and stained with lysotracker Red and Hoechst 33342 were imaged and quantitated for the extent of colocalization of the NBD-cholesterol and lysotracker positive compartments and is depicted as mean % co-localization from multiple fields of three independent experiments. Scale bar = 10µm/ 20µm. D) Intracellular bacterial growth was estimated in Mtb–infected THP1 macrophages left untreated (NT) or treated with (HR alone or HRC (HR in combination with 20 µM citalopram (CIT)) for 3 days. The data are represented as mean log10 values ± SEM from triplicate assays of four independent experiments (N = 4). E) Representative confocal images of THP1 macrophages treated with citalopram (CIT) along with NBD-cholesterol for 24 hours and stained with lysotracker Red, and the colocalization percentage is calculated using ImageJ. Scale bar = 20µm. F) Gene expression levels in macrophages infected with Mtb, followed by treatment with HR alone or with citalopram. The relative expression of the specific gene, relative to GAPDH, is represented as fold change with respect to HR ± SEM of triplicate assays from three independent experiments (N=3). P values less than 0.05, 0.01, 0.005 are represented as *, ** and *** respectively.

To further identify the intracellular niche of the accumulated cholesterol, cells fed with NBD-cholesterol were stained with the lysotracker that labels acidic compartments consistent with the lysosomes. While in the untreated cells, minimal levels of NBD-cholesterol co-localized with lysotracker, the majority of the internalized cholesterol in SRT-treated cells was found to be accumulated in these compartments, indicating a pronounced accumulation of cholesterol in the acidic lysotracker positive compartments corresponding to lysosomes with SRT treatment (Figure 2B, C).

In order to understand if this increase in cholesterol biosynthesis was a generic property of SSRIs, another widely used antidepressant, citalopram, was tested for its ability to control intracellular bacterial growth. Surprisingly, the addition of citalopram to the frontline TB drugs-HR did not affect bacterial control in macrophages, unlike the increased control of infection by HRS in comparison to the antibiotics alone (Figure 2D). Remarkably, unlike SRT, citalopram induced only a modest and heterogeneous increase in lysosomal cholesterol accumulation, with the effect restricted to a small subset of cells, and did not enhance expression of the genes involved in cholesterol biosynthesis (Figure 2E, F).

### Sertraline enhances SREBP2 processing in macrophages

The feedback activation of SREBP2 in response to the absence of cholesterol sensing in the ER on account of lysosomal accumulation is well documented (18, 19). To assess whether SRT activates the signaling cascade, levels of mature SREBP2 (mSREBP2) were measured in macrophages following SRT treatment. Indeed, commensurate with the significant induction of the SREBF2 gene in HRS-treated macrophages, in both the global transcriptomic data and with qRT-PCR analysis, macrophages treated with SRT harbored ∼2-fold higher levels of the cleaved, active form of SREBP2 (mSREBP2) by 24h compared to the untreated cells (Figure 3A-D). The relatively unchanged levels of mSREBP2 in both untreated and citalopram treated cells, lend further support to the importance of this signaling axis in the antibiotic potentiating effect of SRT (Figure 3 E, F).

**Figure 3:**
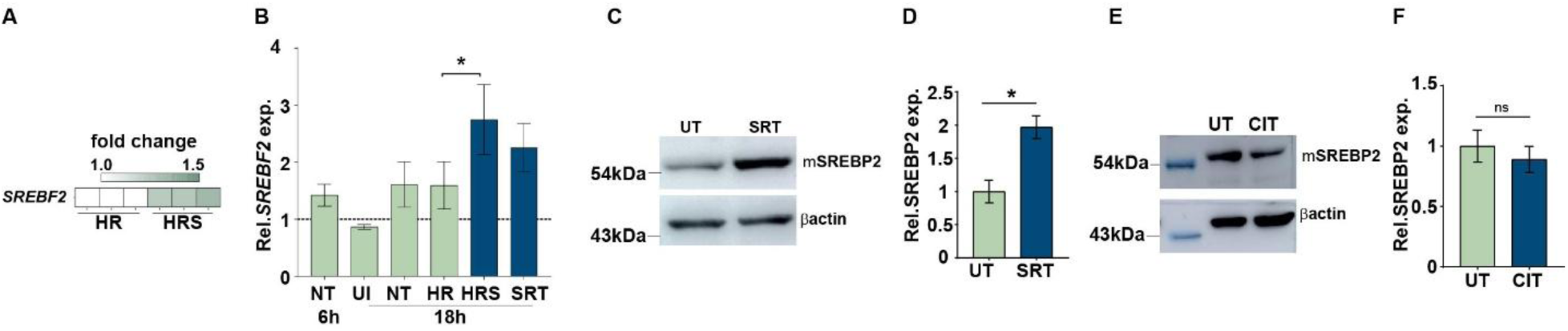
Sertraline enhances SREBP2 processing in macrophages. A -B) Expression levels of *SREB*F2 in the global transcriptomic analysis dataset (A) or quantified by qRT-PCR (B) in Mtb-infected macrophages left untreated (NT) or treated with HR, HRS, or SRT for 18h. The relative expression of specific genes relative to *GAPDH* is represented as fold change + SEM of triplicate assays of three independent experiments (N=3). C-F) The levels of SREBP2 in protein extracts of macrophages left untreated, treated with SRT (C) or 20µM of citalopram (E) for 24 hours were checked by immunoblotting with specific antibodies for SREBP2 and β-actin and quantified by densitometry (D, F). The reactivity for SREBF2 in cells relative to β-actin was quantified in three independent experiments (N=3) and is represented as mean + SEM. P values less than 0.05, 0.01 and 0.005 are represented as *, ** and *** respectively.

### SREBP2 processing is critical for controlling intracellular Mtb growth

SREBP2 activation is regulated by multiple feedback mechanisms; typically, cholesterol and oxysterols have been shown to downregulate this process by modulating the SCAP-SREBP2 egress to the Golgi (20). To test the importance of SREBP2 activation on Mtb control, macrophages were treated with cholesterol exogenously to inhibit SREBP2 processing and tested for their ability to control Mtb growth in the presence of HR or HRS. Addition of cholesterol significantly inhibited SREBP2 processing (2-fold) (Figure 4A, Figure S1A) and resulted in a significant reversal (5-fold) of the effect of SRT on antibiotic potentiation (Figure 4B). Further, the inhibition of SREBP2 activation with a specific S2P protease inhibitor- 1,10-phenanthroline, resulted in a partial but significant reversal of bacterial control by SRT (3-4-fold higher bacterial numbers than HRS alone) (Figure 4C). To further test the relevance of SREBP2 processing, macrophages deficient in the SREBP2 cleavage activation protein (SCAP) were tested for their ability to restrict Mtb growth. As expected, THP1 cells with >50% silencing of *SCAP* with specific shRNAs displayed significantly lower levels of mature SREBP2 in comparison to the cells treated with scrambled shRNAs (SCR) (Figure 4D-F). Consistent with the inhibition of S2P, silencing of SCAP reduced the effectiveness of SRT on Mtb control without affecting the uptake of bacteria (Figure 4G). As observed in Figure 4G, SCAP-silenced macrophages harbored significantly 3-4-fold higher bacterial numbers than the control cells. Even in the untreated cells, loss of SCAP severely impaired the ability to control Mtb with nearly 3-fold higher CFU in the SCAP-deficient cells than in the control (Inset, Figure 4H), corroborating the requirement of SREBP2 maturation in Mtb control by macrophages.

**Figure 4:**
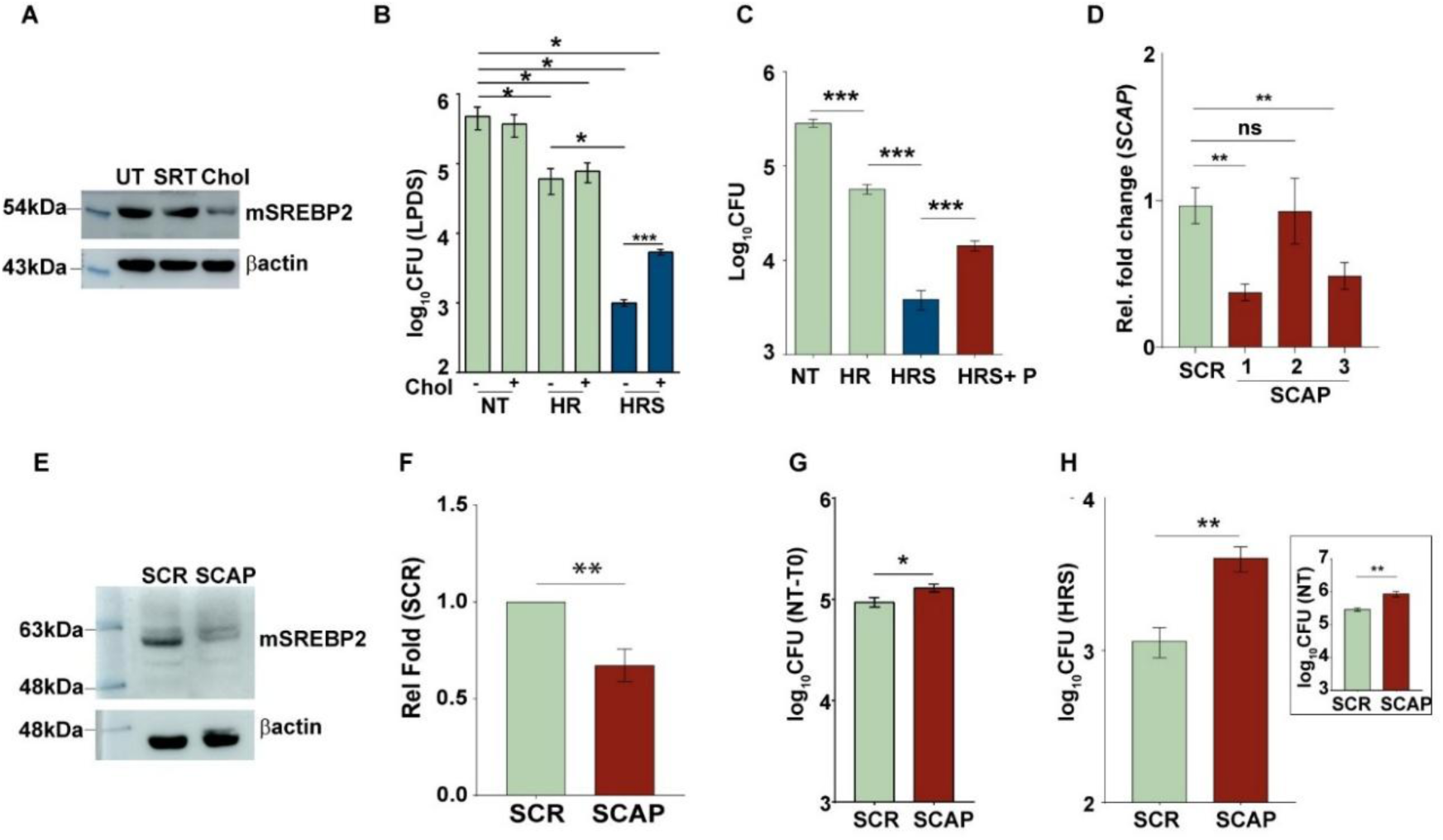
SREBP2 processing is critical for controlling intracellular Mtb growth. A) The levels of SREBP2 in cell extracts of macrophages treated with 20µg/ml of cholesterol for 24 h were analysed by immunoblotting. The levels of β-actin were used as the control values for the comparison. B-C) The intracellular bacterial growth was estimated in Mtb-infected macrophages that were either untreated or treated with (B) cholesterol or (C)- 0.5 mM of 1,10-phenanthroline (P) for 1 hour. This was followed by treatment with HR and HRS alone, in combination with cholesterol, which was replenished every 24 hours in the LPDS media, or with 1,10-phenanthroline. The data is represented as mean log10 values ± SEM from triplicate assays of four independent experiments (N = 4). (D) Evaluation of gene expression levels in *SCAP-silenced* macrophages. The relative expression of the *SCAP* is represented as fold change ± SEM w.r.t. scrambled control in triplicate assays from three independent experiments (N=3). (E) Estimation of mature SREBP2 in protein extracts of *SCAP-silenced* cells by immunoblotting with specific antibodies for SREBP2 and β-actin. The protein marker used is Prestained Protein ladder (MBT092). (F) Densitometric quantification of the reactivity for SREBP2 in cells relative to β-actin of six independent experiments (N=6) is represented as mean + SEM. (G) Bacterial uptake in *SCAP-silenced* cells compared to scrambled control after 6 hours of incubation with Mtb at MOI 5. (H) Intracellular growth of Mtb is depicted in untreated (NT) (inset) or HRS-treated *SCAP-silenced* cells. The data are represented as mean log10 values ± SEM from triplicate assays of three independent experiments (N = 3).

### SRT recapitulates key features of NPC1 inhibition in macrophages

A similar phenotype of organellar sterol accumulation, specifically in the endosomal-lysosomal compartments, is a characteristic feature of NPC1 mutation/ deficiency in the human population (21, 22). A strong binding of SRT and the related antidepressant – indatraline, with the N-terminal domain of NPC-1, has been shown to cause accumulation of cholesterol in lysosomes (23). In order to investigate the relevance of this mechanism, a computational structural analysis using ligand docking for SRT with NPC-1 was performed. The well-established cargo for NPC1-cholesterol, resulted in the identification of a prominent cluster of docked poses with 88% of the docked poses and a mean binding energy of -10.68 kcal/mol (Figure 5A). Having established the correct binding pose, the binding of SRT was tested. Docking of sertraline in the same framework indicated that the docked poses clustered into 2 prominent different clusters, with the top-ranked cluster comprising 56% of the poses, a mean binding energy of -8.28 and -8.13 kcal/mol (Figure 5B). Similar to cholesterol, SRT is also stabilized by hydrophobic interactions and hydrogen bonds, forming a stable protein-ligand complex with a defined pose, indicating its ability to bind to NPC1.

**Figure 5:**
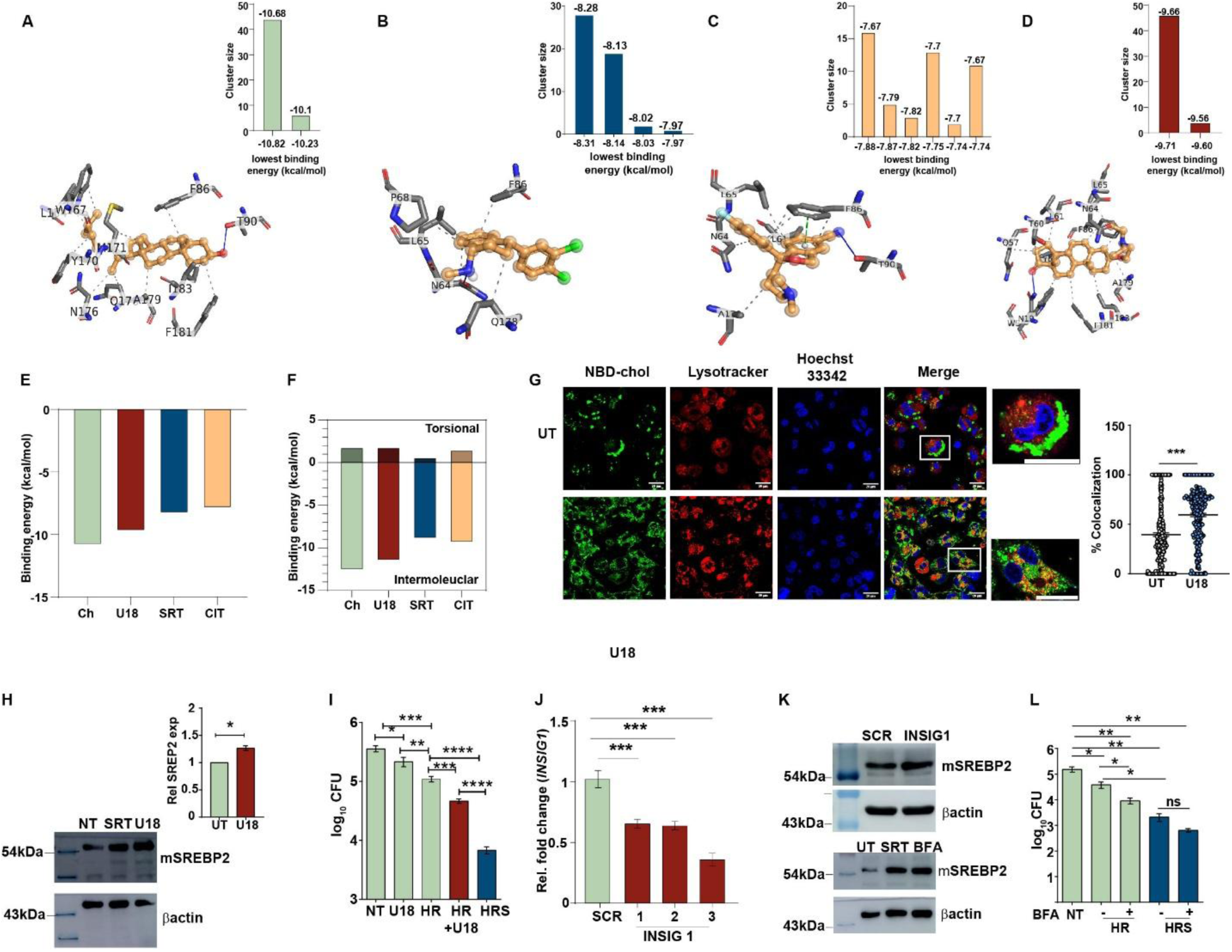
SRT treatment phenocopies NPC1 deficiency. A-D) The best docked poses for Cholesterol (A), Sertraline (B), Citalopram (C), and U18666A (D), having the lowest binding energy. Bar plots represent the clusters of docked poses and binding sites (Mean binding energy (kcal/mol) for each cluster is mentioned at the top of the bar). Grey dotted line: hydrophobic interactions, blue solid line: hydrogen bonds, green dotted line: pi-pi stacking. E-F) The bar plots represent the overall binding energy of each ligand (E), and (F) shows the breakdown of the overall binding energy into intermolecular energy and torsional penalty components. G) The localization of cholesterol was analyzed in cells treated with 2.5 µg/ml of the NPC1 inhibitor- U18666A (U18). The extent of colocalization of NBD-cholesterol and lysosomes (lysotracker) in macrophages was quantified and is expressed as percent co-localization from three independent experiments (N=3) (inset). Scale bar = 20µm. H) The extent of mature SREBP2 was evaluated in macrophages left untreated or treated with U18666A by immunoblotting and densitometric analysis of the lysates. Levels of β-actin expression were used as the control. The relative levels of mSREBP2 in SRT-treated cells (w.r.t. untreated) are represented as relative mean expression + SEM from six independent experiments, as shown in the box (N=6). I) Growth of Mtb in macrophages that were left untreated or treated with HR alone or in combination with U18666A and HRS for 3 days. The CFU numbers are represented as mean log10 values ± SEM from triplicate assays of four independent experiments (N = 4). J) Expression of *INSIG1* gene was quantified by qRT-PCR. *GAPDH* expression values were used as controls for normalization. The relative expression in the *INSIG1* silenced cells was compared with the scrambled control and is represented as Rel. fold change mean ± SEM of triplicate assays from three independent experiments (N=3). K) The levels of mature SREBP2 or β- actin were analyzed by specific immunoblotting of the protein lysates from *INSIG1-silenced* macrophages (Upper panel) and in THP1 macrophages left untreated or treated with 3µg/ml BFA (Lower panel). Levels of β-actin expression were used as the control. L) Growth of Mtb in macrophages left untreated or treated with HR, HRS alone, or in combination with brefeldin A (BFA) for 3 days. The CFU numbers are represented as mean log10 values ± SEM from triplicate assays of three independent experiments (N = 3). P values less than 0.05, 0.01, and 0.005 are represented as *, ** and ***, respectively.

Given its inability to impact lysosomal cholesterol distribution, the other SSRI, citalopram, was evaluated for its ability to bind to NPC-1. In contrast to cholesterol and SRT, the docked poses were found to be distributed across six different clusters, with the top-ranked cluster comprising 32% of the docked poses (mean binding energy = -7.67 kcal/mol), indicating the absence of a defined pose for citalopram in the NPC1 site, suggestive of poor binding to NPC-1 (Figure 5C). To test the relevance of this binding on bacterial control, the specific inhibitor, U18666A, was tested for its NPC-1 binding capacity and the ability to enhance antibiotic efficacy in macrophages. Surprisingly, U18666A also bound specifically to the same binding pocket as cholesterol with a prominent cluster comprising 92% of the docked poses, having a mean binding energy of -9.66 kcal/mol (Figure 5D) (Interactions listed in Suppl table S1). The binding energies computed for the best docked poses of the ligands, cholesterol, and U18666A indicate high affinities in both cases (binding energies -10.82 kcal/mol and -9.71 kcal/mol, respectively), possibly due to the stabilization by a large number of hydrophobic interactions. While the best docked poses of SRT also indicated a high affinity (binding energy –8.31 kcal/mol), citalopram bound NPC1 at a lower affinity of -7.88 kcal/mol (Figure 5E).

In addition, Citalopram had a higher torsional penalty than SRT, lending it a poorer ligand to NPC1 as compared to sertraline (Figure 5F). Further analysis of cholesterol levels in U18666A-treated macrophages revealed a pronounced accumulation in the lysotracker positive compartments with the consequent SREBP2 processing and activation, with an enhanced expression of the cholesterol biosynthetic machinery similar to SRT (Figure 5G, H, S1B). To verify the importance of increased SREBP2 processing in Mtb control, the effect of antibiotics was tested in cells treated with U18666A. Addition of U18666A to the frontline TB drugs -HR resulted in a 3-4-fold better control of Mtb in macrophages by day 3 of infection, emphasizing the relevance of lysosomal cholesterol loading in bacterial control (Figure 5I). To simulate conditions of enhanced cholesterol biosynthesis via increased SREBP2 activation, we resorted to silencing INSIG1, the inhibitor of SCAP-SREBP2 translocation for processing and activation at the Golgi, and observed a significant (>60%) decrease in the levels of INSIG1 expression (Figure 5J). However, we did not observe any detectable change in the levels of mature SREBP2 in these macrophages (Figure S1C). As an alternative, macrophages were treated with brefeldin A (BFA) to facilitate constitutive proteolysis and activation of SREBP2 due to the collapse of the ER-Golgi network(24). In sync with our expectations, the addition of BFA significantly increased the expression of active SREBP2 in macrophages (2-fold- Figure 5K, Figure S1D) that reflected as a 3-4-fold decrease in bacterial numbers in HR treated cells, further emphasizing the importance of SRT-mediated increase in SREBP2 activation in the ability of macrophages to restrict bacteria (Figure 5L).

### Loss of lysosomal integrity couples SRT-mediated cholesterol accumulation with activation of inflammasomes

Accumulation of cholesterol in lysosomes has been associated with a significant damage to the lysosomal membrane, resulting in the formation of pores, a process called lysosomal membrane permeabilization (LMP) (23, 25). This loss of membrane integrity is rapidly repaired by the recruitment of the ESCRT complex via the lectin-galectin 3 (26). Treatment with SRT significantly induced the expression of *LGALS3* in Mtb infected macrophages (Figure 6A-B). To check if SRT compromised the membrane integrity of lysosomes, cells expressing GFP-tagged galectin 3 were treated with SRT. As observed in Figure 6C, galectin3 was distributed uniformly in the untreated cells, in contrast to the punctate signal in the cells treated with LLOme, a dipeptide documented to induce lysosomal damage (27). Interestingly, SRT- treated cells also displayed punctate signals, unlike the treatment with citalopram (Figure 6D), suggestive of extensive lysosomal damage. A close connect between the host lysosomal cholesterol levels and mitochondrial function has been established (28–30). Accumulation of cholesterol in the lysosomes has been associated with altered mitochondrial physiology, exacerbated ROS production, and disorders (31–34). Given our previous observation for an important role for inflammasome activation by SRT (9) via the induction of mitochondrial ROS (10) and an elegant study by Guo et.al, 2018, establishing a direct link between the activation of SREBP2 and inflammasome induction (35), the role of cholesterol accumulation by SRT as a putative cause of inflammasome activation was evaluated in Mtb infections. In line with our previous studies, while SRT induces significant ROS, citalopram failed to activate mitochondrial ROS in macrophages (Figure 6E-G). A direct link between the host cell cholesterol accumulation, increased SREBP2 processing, and inflammasome activity was probed in cells by estimating the levels of IL1β secretion in the different macrophages. Inhibition of SRT-mediated SREBP2 maturation in SCAP-deficient cells reduced the secreted IL1β levels (Figure 6H). This study firmly establishes the importance of the affinity of cationic amphiphilic drugs to the NPC1 protein in augmenting bacterial control, thereby providing a novel target for the future development of host-directed therapy against infections.

**Figure 6:**
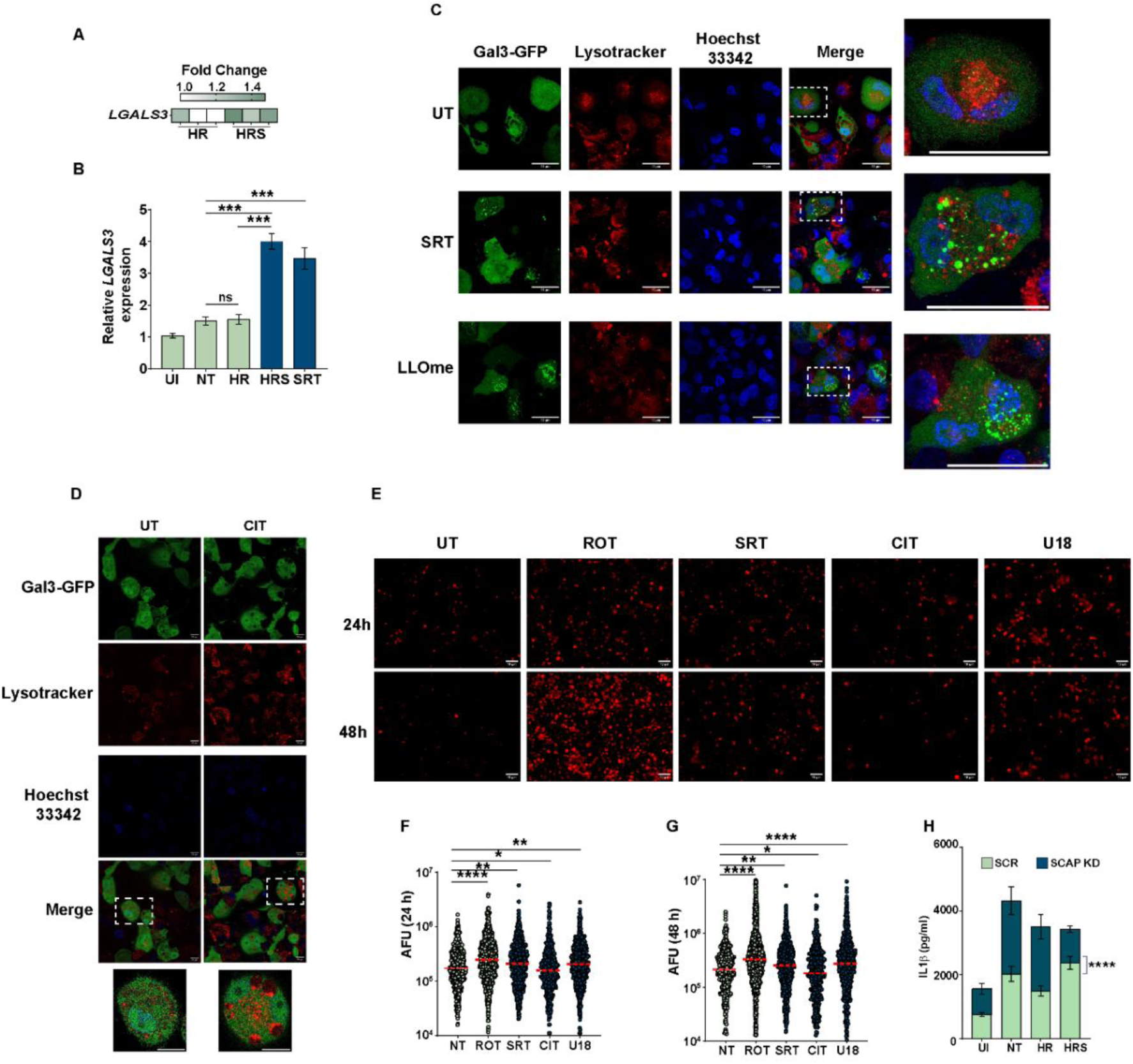
SRT-induced lysosomal cholesterol accumulation induces macrophage inflammasome. (A-B) Gene expression of *LGALS3* (A) in the transcriptomics data and quantified by qRT-PCR (B) in the Mtb-infected macrophages after 18 hours of treatment with HR, HRS, and SRT. The relative expression of specific genes relative to *GAPDH* is represented as fold change + SEM of triplicate assays of three independent experiments (N=3). (C) Representative images of macrophages overexpressing GFP-tagged galectin 3 protein treated with SRT for 24 hours or LLOme for 15 min and stained with lysotracker Red. Boxes indicate insets, magnified on the right. Scale bar = 10µm. (D) Images after citalopram treatment for 24h in GFP-tagged galectin3 overexpressing macrophages. The inset represents the zoom-in. Scale bar = 10µm. (E-G) Representative images showing mitoSox staining after treatment with sertraline (SRT), citalopram (CIT), U18666A (U18), and rotenone (ROT) as the positive control for mitochondrial ROS induction. The extent of ROS was quantified after 24 h (F) and 48 h (G) of treatment and is represented as arbitrary fluorescence units (AFU) from ∼ 800 cells of three independent experiments (N=3). (H) IL-1β levels in the supernatants of Mtb-infected SCR and *SCAP-silenced* cells following treatment with HR or HRS, compared to the untreated group. P values less than 0.05, 0.01 and 0.005 are represented as *, ** and *** respectively.

## Discussion

Infection control by innate immune cells is a complex process with the involvement of multiple modalities, intertwining metabolic and inflammatory response pathways to ensure pathogen removal (36, 37). The complexity only increases with the pathogen adept at modifying signaling cascades and occupying intracellular niches that are designed for infection control (38, 39). The incidence of TB, particularly, has been strongly associated with severe malnutrition(38, 39). Epidemiological data suggest a strong inverse correlation between serum cholesterol levels and TB incidence and mortality, with TB patients displaying severe hypocholesterolemia (40–42). In fact, the enhanced levels of sputum sterilization in a small set of pulmonary TB patients on a cholesterol-rich diet only highlight the importance of improving sterol levels for improved treatment outcomes(43). Recent studies have also implicated a significant role of cholesterol and its derivatives in dictating macrophage response dynamics (44, 45). In line with these observations, we identified an important role for the host cell cholesterol metabolic pathway in the antibiotic-mediated potentiation of Mtb control by SRT.

In this context, we demonstrate that SRT treatment enhances macrophage cholesterol biosynthesis programs through robust activation of the SREBP2 processing, leading to elevated intracellular cholesterol. We reasoned that altering the cholesterol biosynthetic pathway in macrophages would modulate the effects of sertraline. To our surprise, sertraline activity remained unaltered even in conditions of restricted biosynthesis via statins (46). Contrary to our expectations, SRT treatment in the presence of statins increased the expression of the cholesterol biosynthetic genes in macrophages, resulting in enhanced effect of antibiotics in controlling infection, implying the compensatory activation of alternative pathways to maintain homeostasis of cholesterol in cells. Notably, the SREBP2 axis emerged as a critical focal point connecting cholesterol metabolism and infection control in macrophages. In concordance with recent studies that suggest a direct role for statins, we also observed significant activation of SREBP2 with SRT treatment in macrophages is consistent with increased bacterial control (47–49). However, both genetic perturbation (silencing *SCAP*) and pharmacological activation (BFA), but not silencing of *INSIG1* (induced by SRT in macrophages), confirmed the essential role of this pathway in SRT-mediated Mtb restriction.

Our data relates the altered cholesterol trafficking and storage as vital mediators of SRT function. Recently, TMEM97, a cholesterol-responsive NPC1-binding protein, has been implicated in cholesterol trafficking between lysosomes and ER; RNA-interference mediated restraint of TMEM97 reduces lysosomal lipid storage and restores cholesterol levels in the ER in cell models of NPC1 (50). SRT-treated macrophages with increased expression of TMEM97 (Data not shown) resonate with phenotypes of Niemann-Pick type C disease, with cholesterol-laden lysosomes and disrupted ER-lysosome crosstalk. Along similar lines, treatment with the potent NPC1 inhibitor U18666A induced significant dysregulation of cholesterol trafficking from lysosomes to the endoplasmic reticulum (ER). Consistent with this, cells treated with either SRT or U18666A but not with citalopram exhibited markedly enlarged lysosomes, indicative of cholesterol accumulation.

This lysosomal sequestration appears to be a key trigger for downstream immune responses. Oxysterols, particularly 25-hydroxy cholesterol (25HC), a dominant negative regulator of cholesterol biosynthesis (51)acts as a dampener of inflammation, preventing the overstimulation of macrophages by downregulating cholesterol biosynthesis and SREBP2 processing, in addition to activating the esterification of excess free cholesterol (52, 53). 25HC is documented to protect macrophages against mitochondria induced NLRP3 inflammasome activation with ch25h^-/-^macrophages the gene encoding for 25HC, elaborating significantly high levels of IL1β in response to LPS stimulation (54, 55). Mitochondrial ROS, SCAP-SREBP2, and the *FASN* effectively mediate NLRP3-mediated inflammasome activation in cells (35, 56, 57). Reduced expression of *CH25H* (2-fold) and upregulation of *FASN* (∼ 3-fold) suggest a coordinated suppression of anti-inflammatory oxysterol signaling and a shift towards an activated inflammatory state. This metabolic-inflammatory axis distinguishes SRT from related SSRIs such as citalopram, which lacks these effects. In contrast, citalopram had an insignificant effect on these processes, thereby validating the critical role of host cell cholesterol metabolism in macrophage-mediated control of Mtb. These findings further support a model in which host cell cholesterol metabolism plays a critical role in regulating mitochondrial stress and inflammasome activation, thereby contributing to macrophage-mediated control of Mtb.

Further, the observed increase in assembly of galectin 3 close to the lysosomes in SRT-treated macrophages advocates for extensive lysosomal membrane permeabilization (LMP) as a result of lysosomal cholesterol accumulation, which is a property of lysosomotrophic molecules like cationic amphiphilic drugs (CADs) (58). While similar effects were observed with SRT and U18666A, with strong experimental evidence as a CAD, citalopram, a less well-documented cationic amphiphile, failed to bind to NPC-1 protein directly to induce cholesterol accumulation and resultant LMP.

Our findings advance a previously understudied axis of reprogramming macrophage cholesterol metabolism and the consequent induction of a lysosomal stress response that enhances inflammasome activation and bacterial clearance (as seen with SRT). While we effectively establish a direct correlation between the induced LMP and the subsequent activation of mitochondrial ROS, in line with previous reports (59, 60), a detailed analysis of the molecular mechanism by which SRT and U18666A mediates LMP-induced mitochondrial ROS activation will be part of a comprehensive future manuscript. While one can envisage a possible contribution also of accumulated cholesterol in altering the intracellular distribution of frontline TB drugs, leading to enhanced bacterial control, these will be part of future work and communications.

Our current findings thus emphasize the significance of evaluating host metabolic programs for the design of future adjunct therapies for infection control, specifically in the population cohorts with malnutrition and dyslipidemia. Taken together, our work integrates lipid metabolism and innate immune signaling and places host cholesterol metabolism as a crucial determinant of TB pathogenesis and treatment outcome.

### Author contribution

KB and VR were instrumental in the design of the work. KB, RS, NB, JK, MM, and DS were involved in the conduct of the work. SJ and SB were involved in the analysis of the transcriptome high throughput data. None of the authors have any conflict of interest.

## Acknowledgements

The authors thank CSIR (VR- MLP2106) for supporting the study. CSIR-STS0016 is acknowledged for continuous maintenance of BSL3 and ABSL2 facilities. CSIR- BSC0403 is duly acknowledge for the microscopy facility. The student fellowships from CSIR- India (KB) are acknowledged. Biorender.com is duly acknowledged for the graphical abstract preparation. The authors thank Ms. Garima Maurya and Mr. Lakshay Kumar for proofreading and suggestions on improving the manuscript. The authors thank Dr. Michael Glickman for the LPDS media.

## KEY RESOURCE TABLE

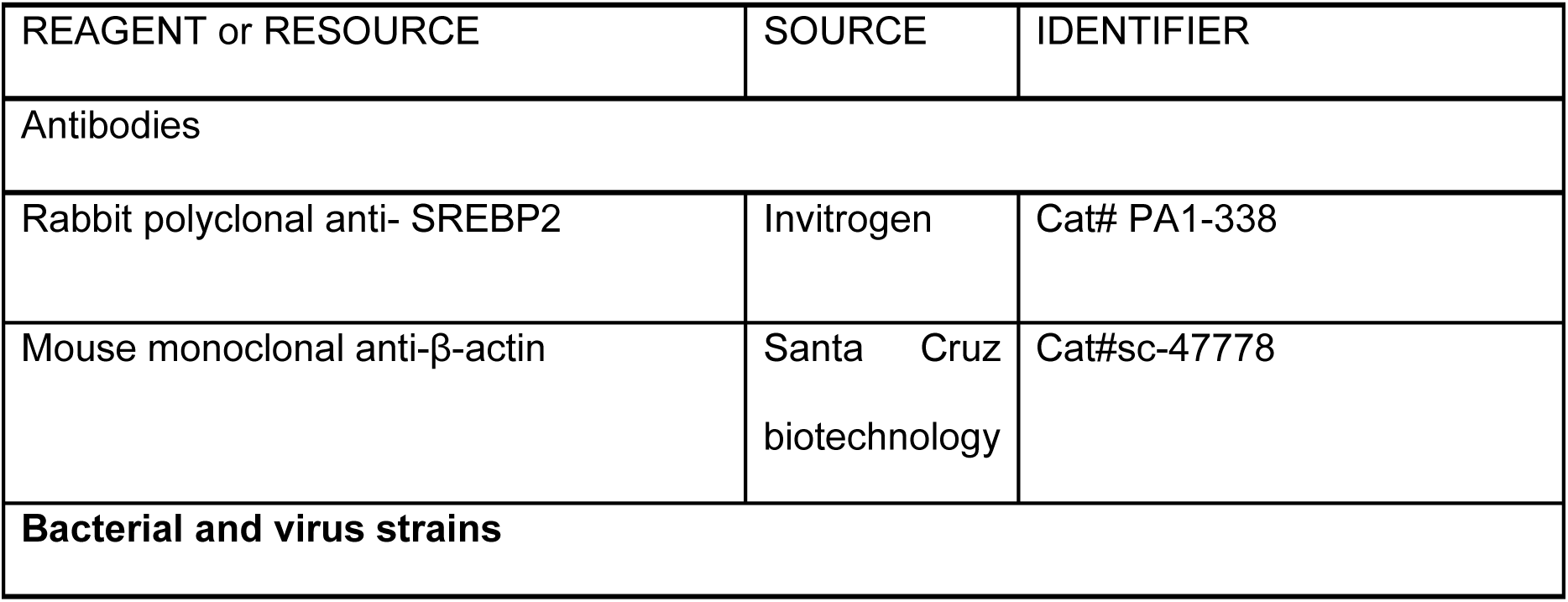
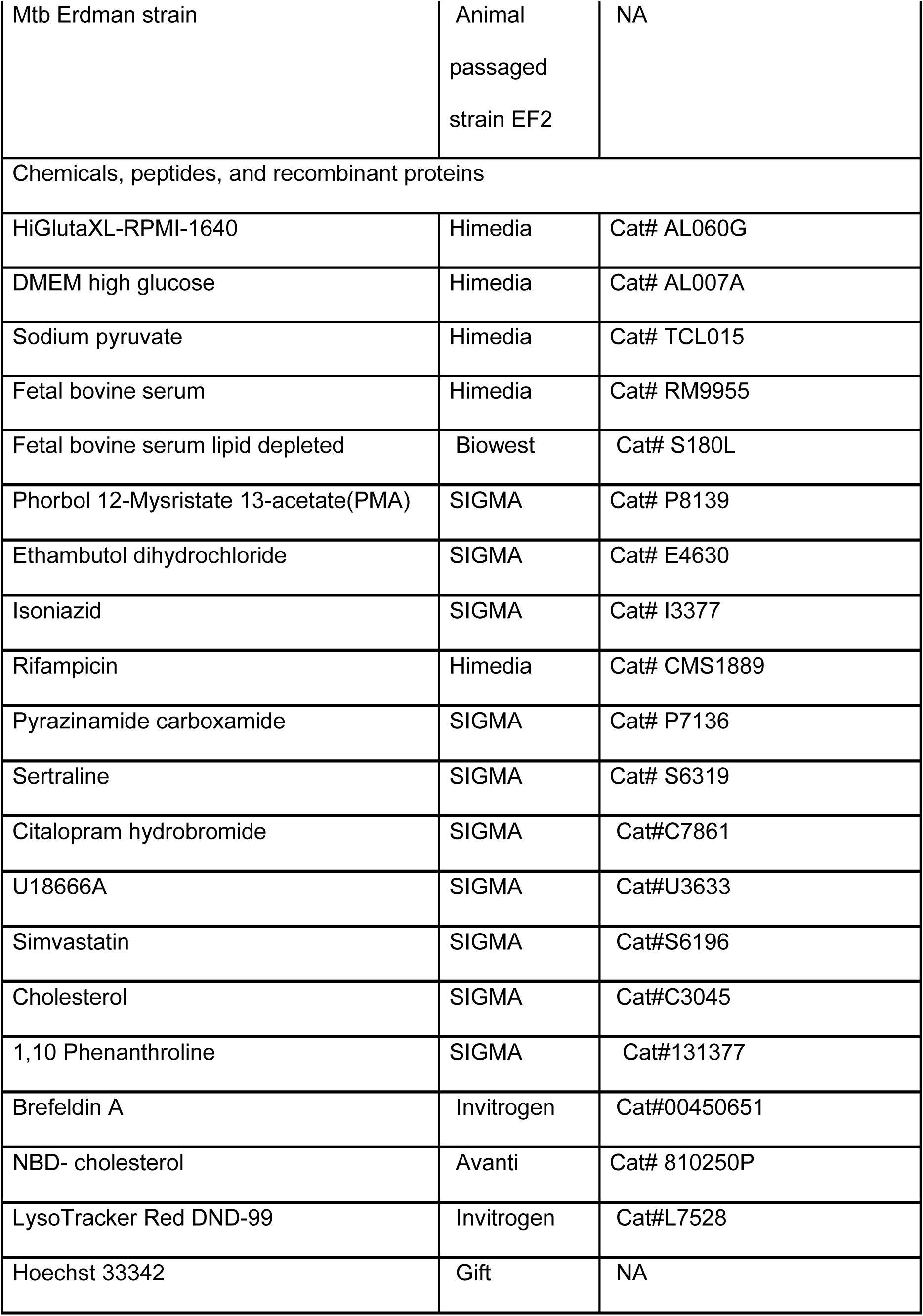
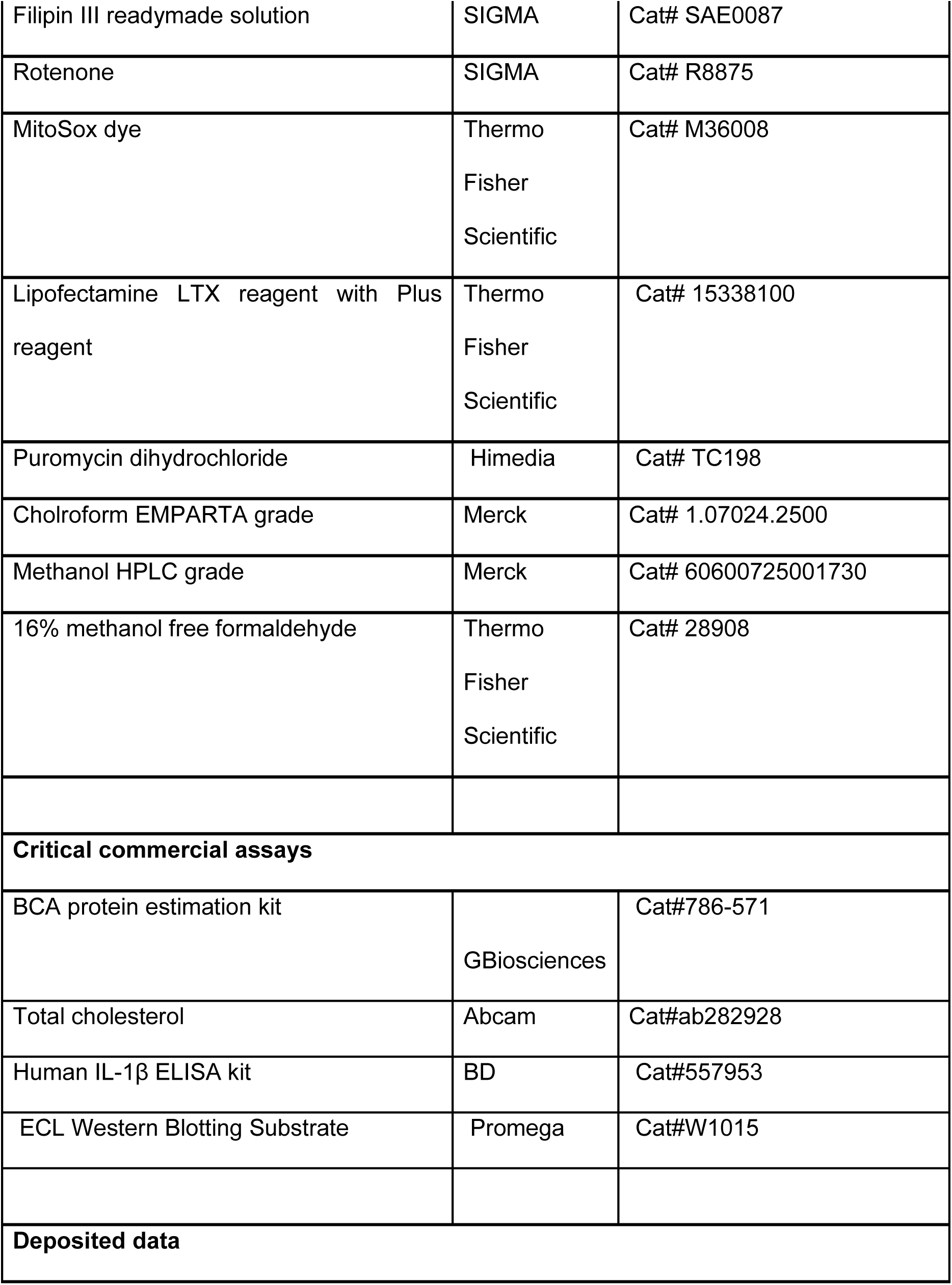
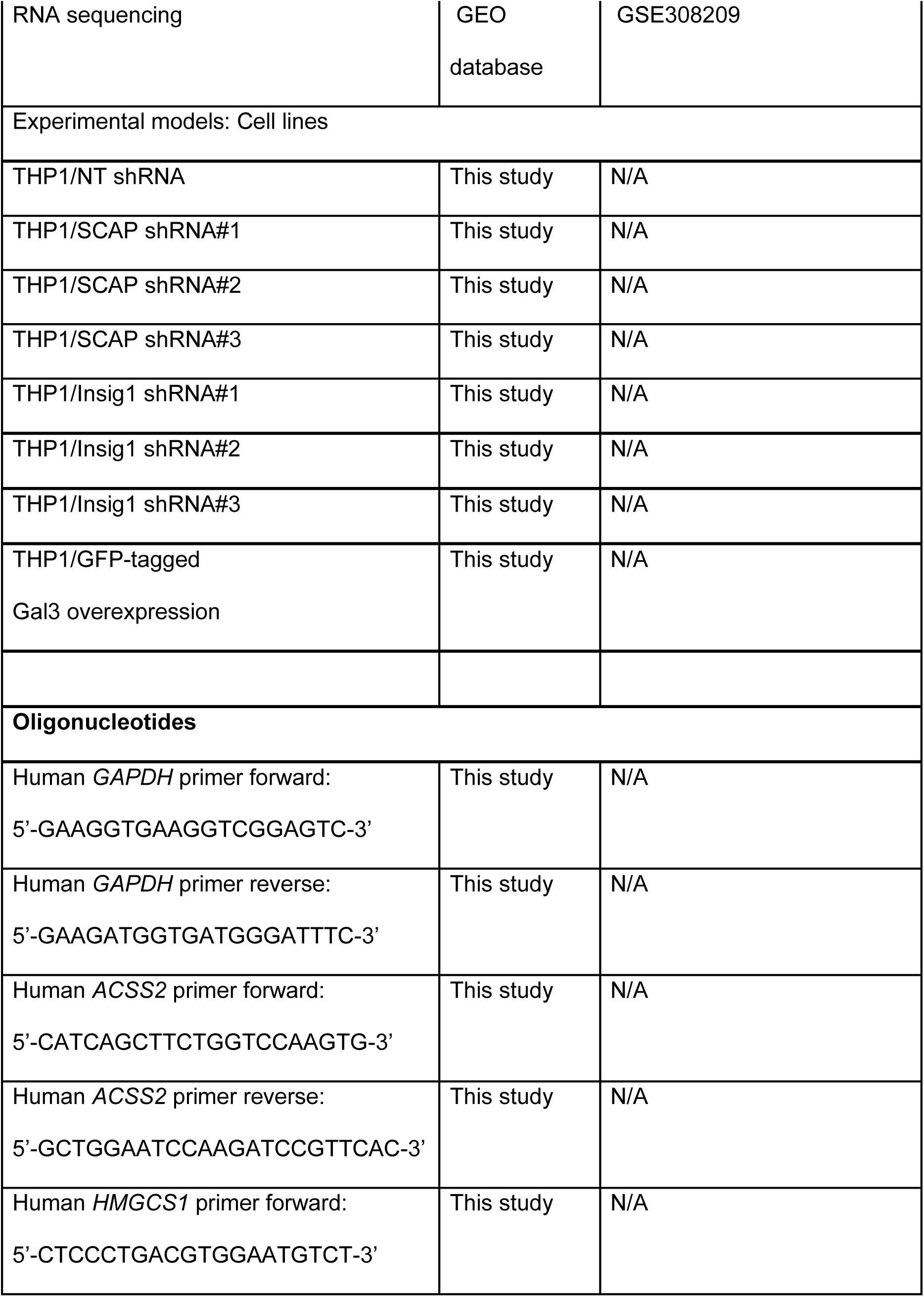
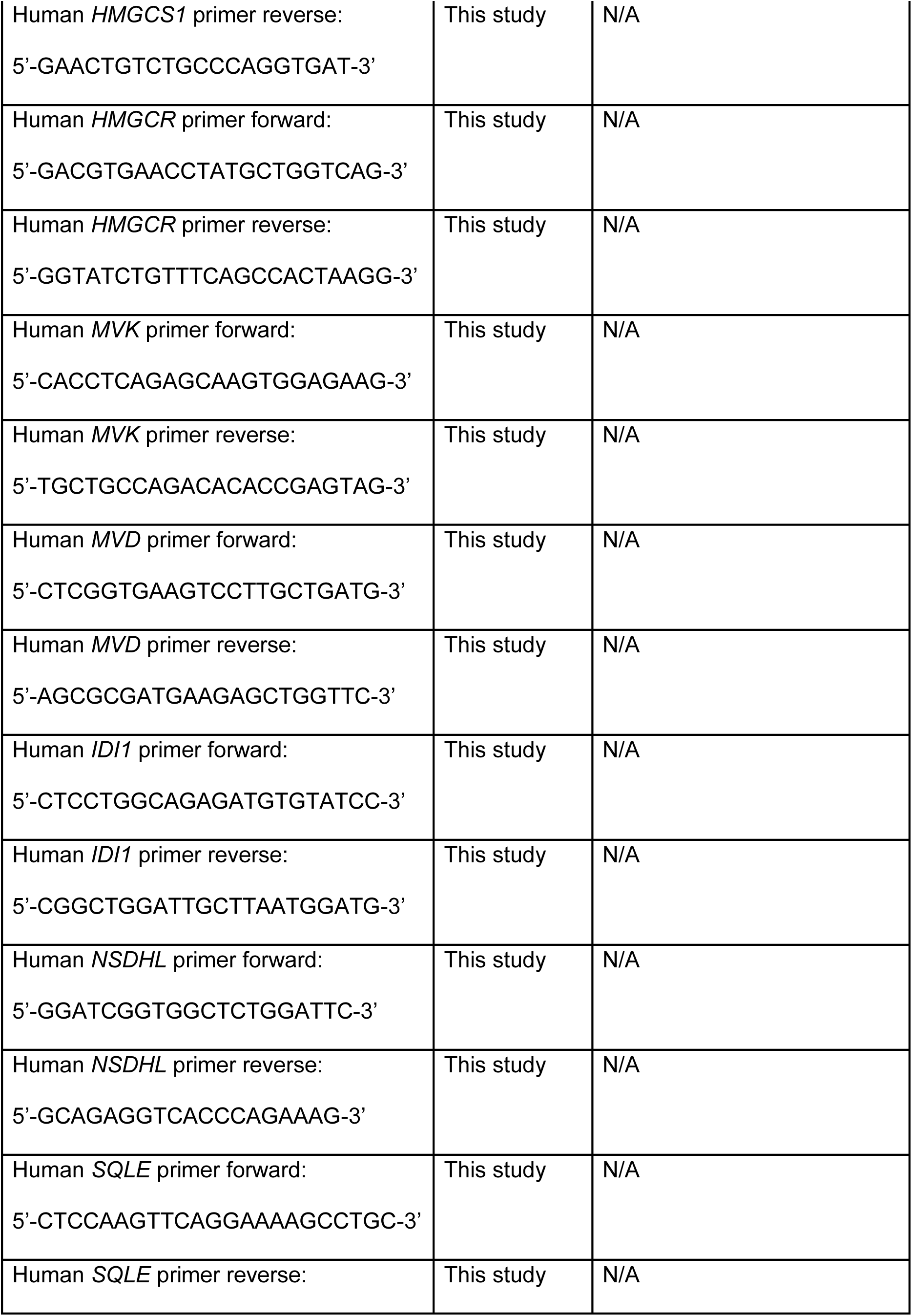
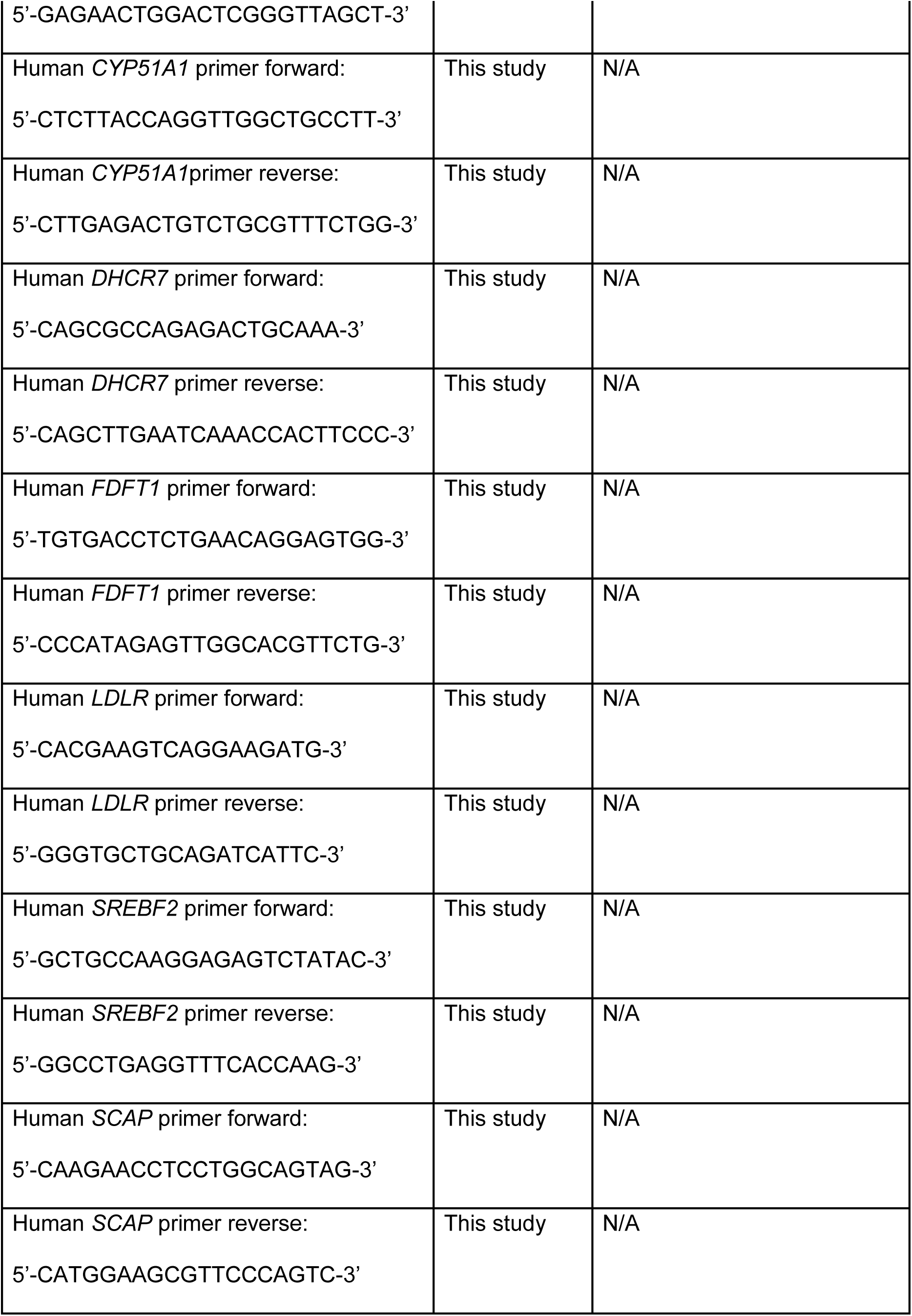
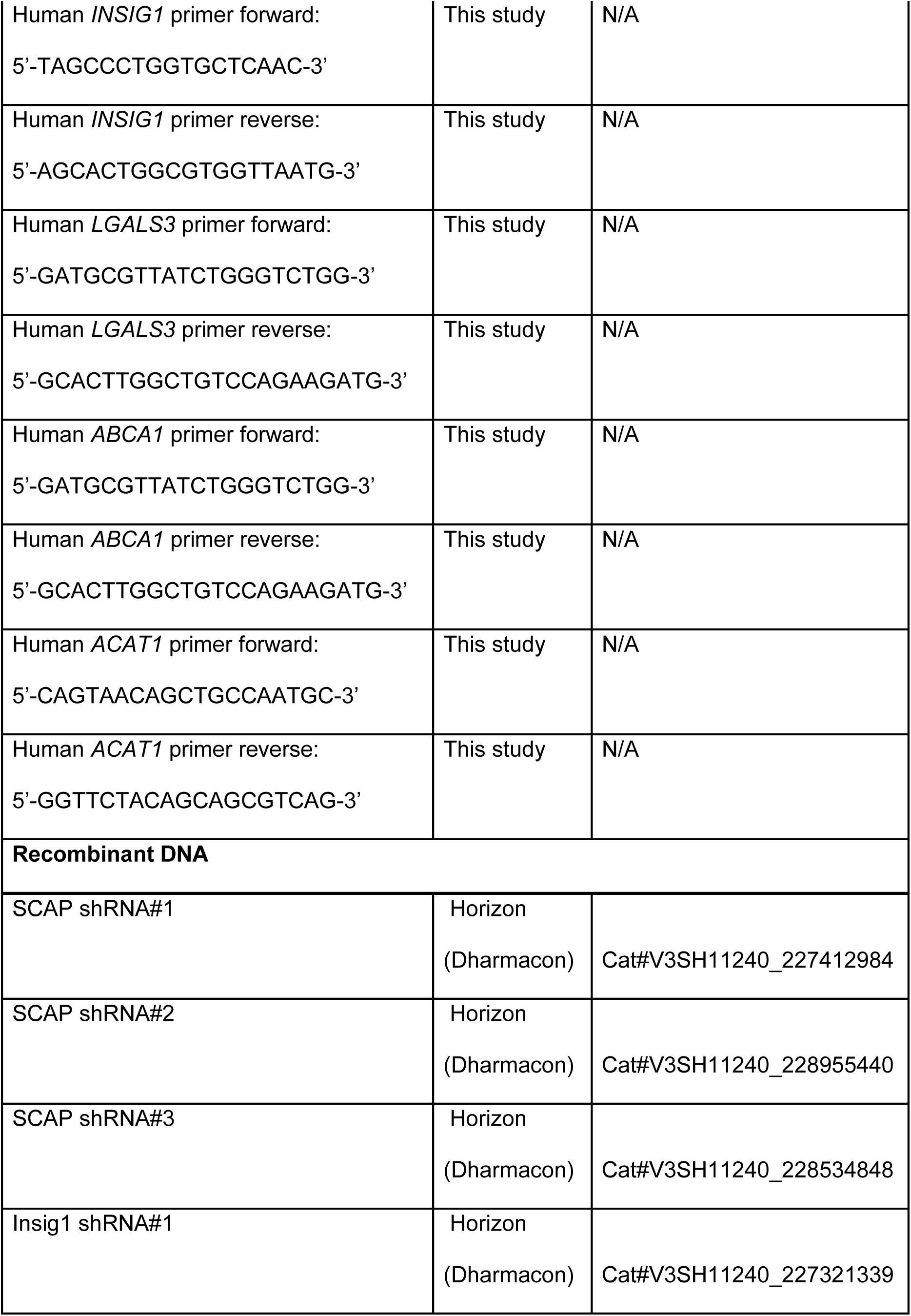
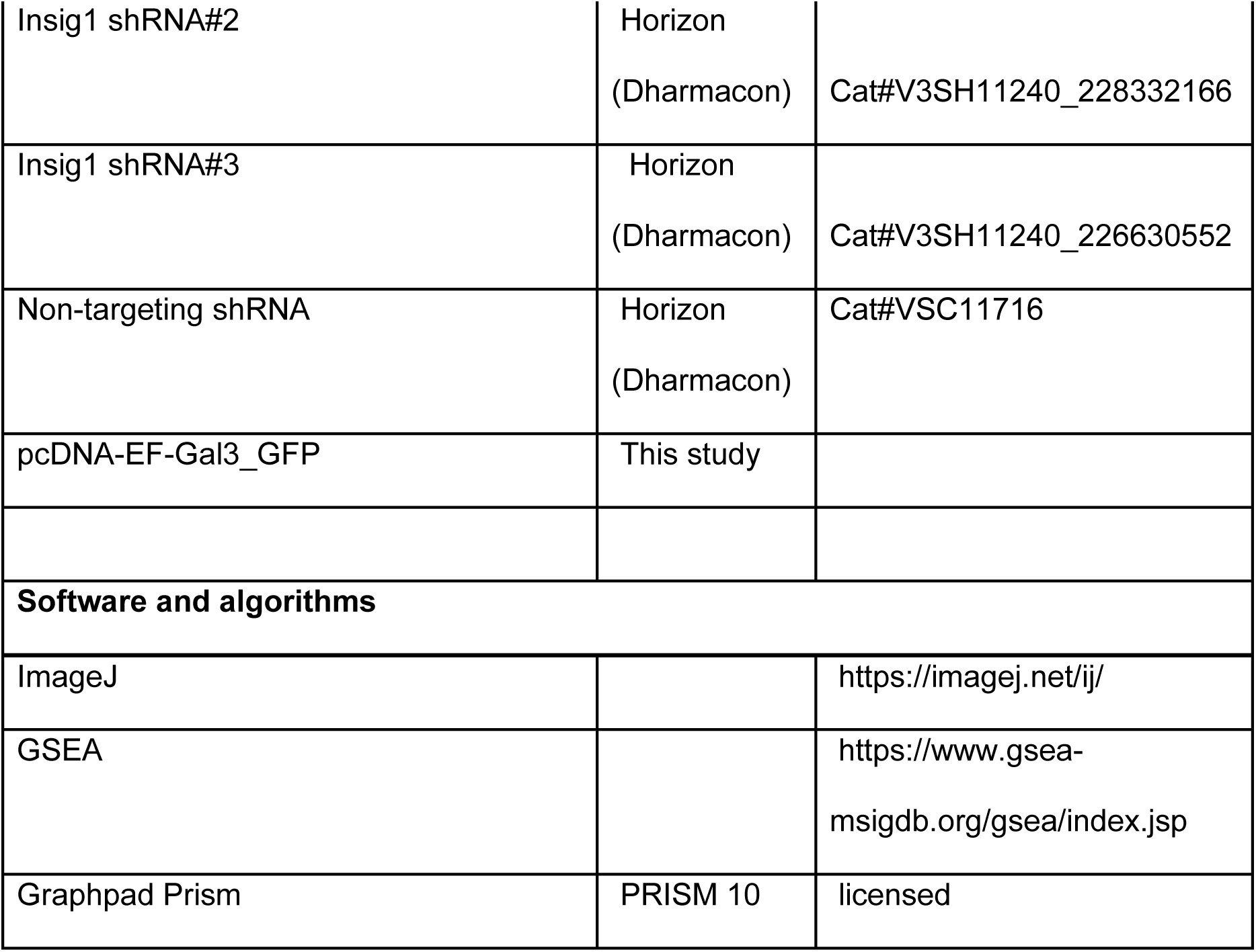

## EXPERIMENTAL MODEL DETAILS

### CELL CULTURE

THP1 and THP1 dual monocytes purchased from ECACC were used in this study. Both THP1 and THP1 dual monocytes were cultured in RPMI 1640 containing 10% FBS and 1 mM sodium pyruvate at 37°C in a 5% CO_2_ humidified incubator. They were differentiated into macrophages using 100 nM Phorbol-13-myristate 13-acetate (PMA) for 24 hours, followed by media change. Following a resting period of 48 hours, cells were utilized for respective experiments.

### BACTERIAL STRAINS AND MEDIA

Mtb Erdmann strain was grown in Middlebrook 7H9 medium with 4% Albumin-dextrose-saline Or Middlebrook 7H10 plates supplemented with 10% oleic acid-albumin-dextrose-catalase (OADC) at 37°C. A final concentration of 0.05% Tween 80 was added to the liquid medium.

## METHOD DETAILS

### RNA sequencing and analysis

Total RNA was isolated from the Mtb-infected macrophages after treatment with anti-TB drugs (Isoniazid and rifampicin) in the presence or absence of sertraline for 18 hours. Total RNA was used for stranded library preparation and paired-end clustering with 150bp x 2 end of sequencing performed on HiSeqX (Illumina) by Medgenome Labs. Raw and processed RNA-Seq data are openly available on GEO database (GSE308209). Quality check of raw sequenced reads was performed using Fastqc 0.12.1 and summarized with Multiqc 0.4. Adapters (Illumina universal adapters) and poor-quality bases were trimmed using Trimmomatic 0.39. The trimmed reads were aligned to the human genome gtf (v44) using Star (2.7.10b). Finally, the rawcounts matrix was generated using Subread 2.0.6. Differential gene expression analysis was conducted via edgeR3.42.4. Gene set enrichment analysis was used to compare datasets.

### Quantitative PCR

RNA was isolated using RNAzol (Sigma), and cDNA was synthesized using oligo-dT primers using Verso cDNA synthesis kit following the manufacturer’s protocol. Quantitative PCR was performed using KAPA SYBR green (Sigma) on a LightCycler480 (Roche).

### Total cholesterol estimation

THP1 monocytes were seeded in a 24-well plate at a density of 0.5*10^6 cells. Cells after washing with 1X PBS were lysed in 100ul of 1% Triton X-100 for 20 mins on ice. Protein was estimated by the BCA protein estimation kit following the protocol given. For total lipid isolation, 400µL of chloroform: methanol (1:2) was added along with 100µL of 50 mM citric acid. The mix was vortexed and spun down. After the addition of 100µL of autoclaved water, the mix was vortexed and spun down, and finally, 100µL of chloroform was added. After centrifugation of the mix at 12000g for 10 minutes, the lower phase was collected and vacuum centrifuged for 30 minutes. The pellet was diluted in cholesterol assay buffer, and total cholesterol levels were assessed using the Total Cholesterol & Cholesteryl Ester Colorimetric Assay Kit (ab282928) following the recommended protocol. Values were normalized with the protein content.

### Macrophage infection

The Mtb culture was centrifuged and washed with PBST (1x PBS with 0.05% of Tween 80) twice, and a single cell suspension was made after spinning down the culture at 800g for 10 minutes. THP1 dual macrophages were infected for 6 hours at MOI5. After 6 hours, cells were washed twice with 1X PBS, and appropriate cell culture media was added.

### Intracellular bacterial growth assay

THP1 dual monocytes were seeded in a 48-well plate at a density of 0.15 million cells. Cells were differentiated into macrophages as indicated, and after a 48-hour resting period, cells were infected with Mtb for 6 hours at MOI5. After stopping the infection, cells were treated with drugs isoniazid (20ng/mL), rifampicin (100ng/mL), and sertraline (20µM) given in 10% FBS RPMI media. At day 3 post-treatment, cells were lysed with water containing 0.05% Tween 80, and dilutions of intracellular bacteria were made in PBST. The final diluted lysate was plated on 7H10 plates.

To restrict cholesterol uptake by the macrophages, treatment with TB drugs was given in RPMI media supplemented with 10% Lipoprotein-deficient serum (LPDS).

For the cholesterol replenishment assay, Mtb-infected THP1 macrophages were treated with cholesterol (2µg/ml) for an hour after infection, followed by incubation with anti-TB drugs in LPDS medium, and further cholesterol was added every 24 hours.

### Immunoblotting

THP1 dual macrophages were lysed in RIPA buffer along with a protease inhibitor cocktail. Protein concentration was estimated by BCA assay, and 50µg of total protein was loaded per sample in a 10% polyacrylamide gel. Protein was transferred to a nitrocellulose membrane, and the membrane was blocked using a 5% BSA solution. The membrane was next incubated overnight in anti-rabbit SREBP2 antibody, on shaking at 4°C. Next, the membrane was washed thrice with PBST (1X PBS with 0.05% Tween 20) solution and incubated with HRP-conjugated secondary antibody for an hour at room temperature. After washing with PBST, the membrane was imaged using ImageQuant LAS 500. The protein markers used are Prestained Protein ladder (MBT092) and puregene three color prestained protein ladder (PG- PMT2922).

### Filipin Staining

The THP1 macrophages were seeded in confocal chamber slides (1.5*10^4 cells per chamber) and treated with sertraline at 20µM for 24 hours. Cells were washed twice with 1x PBS before fixation with 4% methanol-free formaldehyde in 1X PHEM buffer for 20 mins at room temperature. After an additional wash with 1x PBS, cells were stained with filipin staining solution (50ug/ml diluted in 1x PBS) for an hour in the dark at room temperature. Next, the cells were washed with 1x PBS and imaged on a Leica SP8 laser scanning confocal microscope. Filipin was excited at the 405nm laser.

### Cholesterol compartmentalization assay

THP1 monocytes were seeded at a density of 0.15 million cells in a four-chamber confocal dish and differentiated as mentioned. The NBD-cholesterol at a final conc of 10µM was added for 24 hours along with the different compounds such as sertraline (20µM), citalopram (20µM), and U18666A (2.5µg/ml). Before initiation of imaging, cells were stained with Lysotracker Red (100nM) along with Hoechst dye for 1 hour in 10% FBS containing RPMI. Images were taken using a Leica SP8 laser scanning confocal microscope.

### Stable knockdown/overexpression cell line generation

Lenti-X 293T cells were transfected with pVSVG and pDR8.2 plasmids together with lentiviral constructs expressing gene-specific shRNA or non-targeting controls to generate viral particles. After 6 hours of transfection, the media was changed to 2% FBS containing DMEM, and after 48 hours, the cell supernatant was collected and centrifuged at 100g for 5 minutes to remove cell debris. Further, this supernatant was filtered through a 0.45µ syringe filter, and viral particles were concentrated using a 30kDa centricon filter. For transduction in THP1 monocytes, 50,000 cells were seeded in a 24-well plate, and 100µL of viral stock was added along with a final concentration of 8µg/mL polybrene. After 24 hours media was changed, and antibiotic selection was initiated after 48 hours of transduction with 0.6µg/mL of puromycin. Cells showing tRFP fluorescence co-expressed with shRNA were checked regularly.

The hGal3 and eGFP are excised from the pEGFP-hGal3 plasmid using Nhe1, Nde1, and Msc1 restriction enzymes and ligated to Xba1 and Msc1 digested pcDNA 3.1 vector. The final plasmid is transfected into THP1 monocytes using Lipofectamine LTX.

### Computational assessment of NPC1- SSRI interactions

A composite structure of NPC- 1 protein (cryo-EM, PDB IDL 6W5V) with the addition of missing atoms (PDB Fixer) (61) and validated for structure quality by Procheck (62) for Ramachandran parameters were used as a template for the analysis of interaction with SSRIs- Sertraline and citalopram. Ligand structures of Cholesterol, U18666A, sertraline, and citalopram from the PDB ligand library were converted from SDF to PDB format using OpenBabel (63)and used for docking studies focused on the binding region reported earlier (23). The residues reported to interact with sertraline were selected as the binding site region, with the centroid of the binding site used as the center of the grid box with dimensions (40,40,40 npts) for docking analysis with 50 iterations for each ligand by Autodock4.2 (64). The docked poses were clustered at the default value of 2.0Å RMSD, and the clusters were ranked based on the lowest binding energy in each cluster. The binding energy is calculated in AutoDock as a semi-empirical function of intermolecular forces such as Vanderwaals forces, hydrogen bonds, desolvation energy, electrostatic energy, and torsional energies (65). The binding energy of the best docked pose, the mean binding energy of the top cluster, and the total number of clusters were noted. The docked pose with the lowest binding energy was visualised using PyMOL, and the protein-ligand interactions were predicted using PLIP (66).

### Galectin 3 puncta assessment

GFP tagged galectin-3 overexpressing THP1 monocytes were seeded in a chambered confocal dish and treated with sertraline (20µM) and citalopram (20µM) for 24 hours, and with 1mM LLOMe for 15 minutes. Before imaging, cells were stained with Lyostracker Red and Hoechst for an hour. Images were taken using a Leica SP8 laser scanning confocal microscope.

### Mitochondrial ROS estimation

THP1 dual cells were differentiated as indicated. Treatment was given with rotenone at 10µM for 30 minutes or with different indicated compounds. The cells were incubated with 5µM of MitoSox dye for 30 minutes. After washing with 1X PBS, cells were imaged in the Invitrogen™ EVOS™ M5000 Imaging System.

### ELISA

THP1 dual macrophages were infected with Mtb at MOI5 and treated with different compounds. Supernatant was collected at the indicated time points and filtered through a 0.2µ filter. Concentration of IL-1β was determined using an ELISA kit following the recommended protocol.

### Statistical analysis

The results are depicted as mean ± SEM. Statistical analysis was carried out using Student’s t-test (Welch’s test or Mann-Whitney test) using GraphPad Prism. P values less than 0.05, 0.01, 0.005 are represented as *, ** and *** respectively.

---

**Fig S1.**
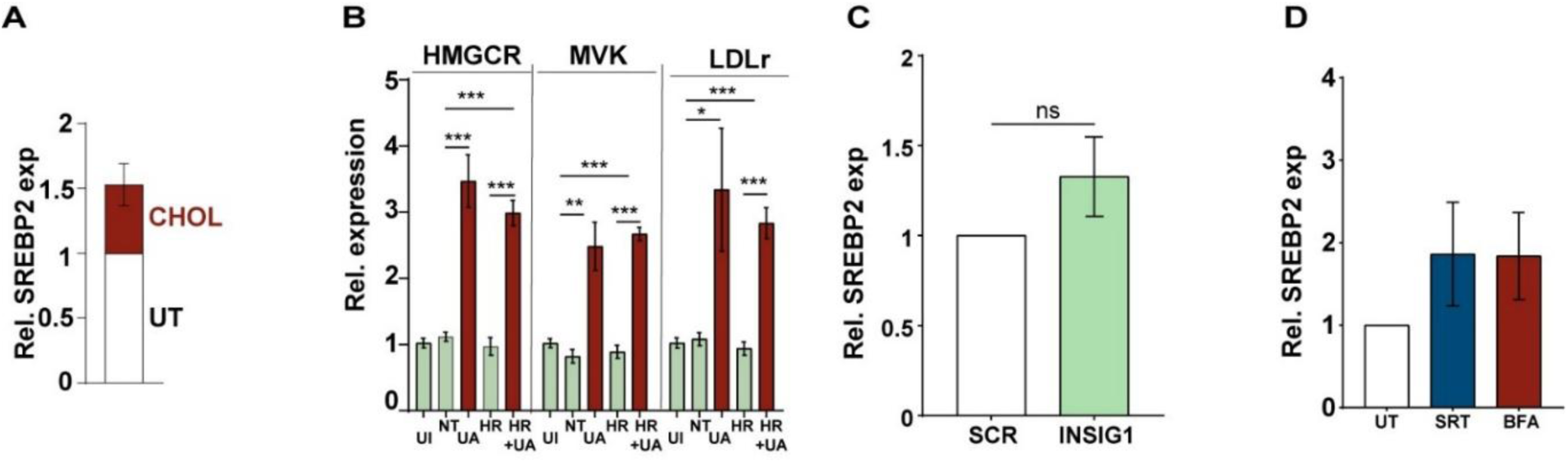
A) The extent of SREBP2 activation was evaluated in macrophages left untreated or treated with cholesterol for 24h by immunoblotting and densitometric analysis of the lysates. Levels of β- actin expression were used as the control. The relative levels of mSREBP2 in cholesterol-treated cells (w.r.t. untreated) are represented as relative mean expression + SEM from three independent experiments, is shown in the box (N=3). B) Expression of genes for cholesterol biosynthesis was quantified by qRT-PCR in the Mtb-infected macrophages left untreated or after 18 hours of treatment with HR with and without U8666A (UA). The relative expression of specific genes relative to GAPDH is represented as fold change + SEM of triplicate assays of three independent experiments (N=3). C, D) The extent of SREBP2 activation was evaluated in SCR or *INSIG1-silenced* macrophages (C) or left untreated or treated with 3µg/ml brefeldin A (D) for 24h by immunoblotting and densitometric analysis of the lysates. Levels of β-actin expression were used as the control. The relative levels of mSREBP2 in cells (w.r.t. untreated) are represented as relative mean expression + SEM from three independent experiments, as shown in the box (N=3).

**Table S1:** Binding interactions predicted for the best docked poses of each ligand.

| Cholesterol |  |  |
| --- | --- | --- |
| Amino acid | Ligand atom | Distance in Å |
| **Hydrophobic interactions** |  |  |
| PHE86 | C19 | 3.73 |
| L154 | C25 | 3.64 |
| W167 | C27 | 3.68 |
| Y170 | C24 | 3.98 |
| M171 | C18 | 3.35 |
| N176 | C21 | 3.67 |
| Q178 | C21 | 3.51 |
| A179 | C12 | 3.18 |
| F181 | C2 | 3.95 |
| I183 | C2 | 3.33 |
| **Hydrogen bonds** |  |  |
| T90 | O1 | 2.80 |
| U18666A |  |  |
| Amino acid | Ligand atom | Distance in Å |
| **Hydrophobic interactions** |  |  |
| W5 | C13 | 3.86 |
| Q57 | C16 | 3.16 |
| T60 | C16 | 3.54 |
| L61 | C15 | 3.51 |
| N64 | C23 | 3.88 |
| L65 | C23 | 3.77 |
| F86 | C23 | 3.48 |
| F86 | C3 | 3.34 |
| F86 | C7 | 3.59 |
| T90 | C11 | 3.37 |
| A179 | C25 | 3.99 |
| F181 | C14 | 3.91 |
| I183 | C6 | 3.27 |
| **Hydrogen bonds** |  |  |
| N19 | O2 | 1.86 |
| Sertraline |  |  |
| **Amino acid** | **Ligand atom** | **Distance in Å** |
| **Hydrophobic interactions** |  |  |
| N64 | C8 | 3.73 |
| L65 | C8 | 3.13 |
| P68 | C9 | 3.26 |
| F86 | C7 | 3.08 |
| Q178 | C3 | 3.74 |
| **Hydrogen bonds** |  |  |
| A64 | N11 | 1.99 |
| Citalopram |  |  |

| Amino acid | Ligand atom | Distance in Å |
| --- | --- | --- |
| **Hydrophobic interactions** |  |  |
| L61 | C17 | 3.30 |
| N64 | C15 | 3.85 |
| L65 | C13 | 3.81 |
| F86 | C18 | 3.23 |
| F86 | C10 | 3.56 |
| F86 | C15 | 3.34 |
| A179 | C09 | 3.52 |
| **Hydrogen bonds** |  |  |
| T90 | N04 | 2.35 |
| **pi-pi stacking** |  |  |
| F86 |  | 3.95 |

